# Rbfox splicing factors maintain skeletal muscle mass by regulating calpain3 and proteostasis

**DOI:** 10.1101/257261

**Authors:** Ravi K. Singh, Arseniy M. Kolonin, Marta L. Fiorotto, Thomas A. Cooper

## Abstract

Alternative splicing promotes proteomic diversity important for cellular differentiation and cell fate determination. Here, we show that deletion of the highly conserved Rbfox1 and Rbfox2 alternative splicing regulators in adult mouse skeletal muscle causes rapid, severe loss of muscle mass. Homeostasis of skeletal muscle tissue requires a dynamic balance between protein synthesis and degradation (proteostasis) but the mechanisms that regulate this balance are not well understood. Rbfox deletion did not cause reduced global protein synthesis, but resulted in reduced autophagy flux and altered splicing of hundreds of transcripts including Capn3, which produced an active form of calpain3 protease. The results indicate Rbfox proteins regulate proteostasis in skeletal muscle tissue by control of calpain and autophagy-lysosome pathways.

**Highlights:**

- Proteostasis in adult skeletal muscle is post-transcriptionally regulated, in part by alternative splicing via Rbfox1/2
- Rbfox1/2 regulate hundreds of targets in skeletal muscle, including Calpn3, to maintain muscle mass in adult mice
- Autophagy flux is markedly decreased in muscle lacking Rbfox1/2
- As for neurons, altered proteostasis is detrimental to adult muscle

## INTRODUCTION

Although much research in the muscle field has been devoted to understanding myogenesis, muscle regeneration and myofiber function (Bassel-Duby and Olson, 2006; Comai and Tajbakhsh, 2014; Potthoff and Olson, 2007; Schiaffino et al., 2013; Yin et al., 2013), there has been growing recognition of the importance of maintenance of muscle mass in adulthood. Skeletal muscle plays a crucial role in metabolism (James et al., 2017; Rai and Demontis, 2016) and directly influences quality of life in aging and in chronic disease (McLeod et al., 2016). Muscles also face special challenges: they must be able to generate considerable force on demand and are continually exposed to mechanical, temperature, and oxidative stresses. Maintaining homeostasis under such diverse conditions requires tight regulation of protein synthesis and turnover (Bell et al., 2016b); failure of proteostasis occurs in many muscle diseases (Lecker et al., 2006; Nishino et al., 2000; Richard et al., 1995; Sandri et al., 2013), as well as the more general conditions of sarcopenia and cachexia (Bowen et al., 2015). Despite the common occurrence of these latter conditions, however, we still do not fully understand the mechanisms that are critical for the maintenance of adult muscle mass.

We hypothesized that alternative splicing might perform such a function. Alternative splicing generates multiple protein isoforms from a single gene, thereby expanding the functional repertoire of proteins far beyond what could be expected from the relatively modest size of the genome (Kalsotra and Cooper, 2011; Wang et al., 2008; Yang et al., 2016). The resulting protein diversity is thought to be particularly important in tissues that must respond to highly variable conditions, such as the brain, heart, and skeletal muscle (Kalsotra and Cooper, 2011; Raj and Blencowe, 2015). Indeed, skeletal muscle has particularly high levels of alternative splicing. We and others have shown that the splicing factors Rbfox1 and Rbfox2, which are highly conserved from *C. elegans* to humans (Gallagher et al., 2011; Jin et al., 2003; Kuroyanagi et al., 2007; Venables et al., 2012), are required for muscle differentiation and function (Pedrotti et al., 2015; Runfola et al., 2015; Singh et al., 2014), but Rbfox factors have never been studied specifically in adult animals.

In this study, we induced skeletal muscle-specific knockout of both Rbfox1 and Rbfox2 in adult mice (seven weeks of age or older). Our double knockout mice suffered a rapid, severe loss of skeletal muscle mass (30–50% within four weeks). RNA sequencing of control and double knockout (DKO) skeletal muscles, two weeks after knockout induction, revealed that hundreds of transcripts were altered in expression and splicing. This loss of mass was attributable to an increase in proteolysis rather than a reduction in protein synthesis.

## RESULTS

### Deletion of Rbfox1 and Rbfox2 in adult mice causes rapid, severe muscle loss

To determine the role of Rbfox splicing factors in mature adult myofibers, we induced knockout of Rbfox1, Rbfox2, or both in seven-week-old mice using a tetracycline-inducible and skeletal muscle-specific Cre line (Rao and Monks, 2009). To quantify the knockout efficiency of Rbfox1 and/or Rbfox2, we performed western blot analysis using tibialis anterior (TA) muscle protein extracts from control (Rbfox1 f/f, Rbfox2 f/f), Rbfox1 knockout (Rbfox1^f/f^: *ACTA1-rtTA^cre/+^*), Rbfox2 knockout (Rbfox2^f/f^: *ACTA1-rtTA^cre/+^*), and double knockout (Rbfox1^f/f^: Rbfox2^f/f^ *ACTA1-rtTA^cre/+^*) animals (**Fig. 1A**). Rbfox1-only knockout led to a two-fold upregulation of Rbfox2 protein in muscle, whereas Rbfox1 protein level did not increase significantly in Rbfox2-only knockout muscle. We also tested the effects of Rbfox1 and/or Rbfox2 deletions on alternative splicing of Bin1 exon 10, which we identified previously as an Rbfox target in a myoblast culture cell line (Singh et al., 2014). Single knockout of either Rbfox1 or Rbfox2 diminished the inclusion of Bin1 exon 10 by <5%, but Rbfox DKO reduced inclusion of this exon by >65% (**Fig. 1A**, bottom panel). These observations indicate that Rbfox1 and Rbfox2 paralogs compensate for one another’s loss of function and regulate overlapping splicing events, so the experiments described here were conducted primarily with the DKO animals.

**Figure 1.**
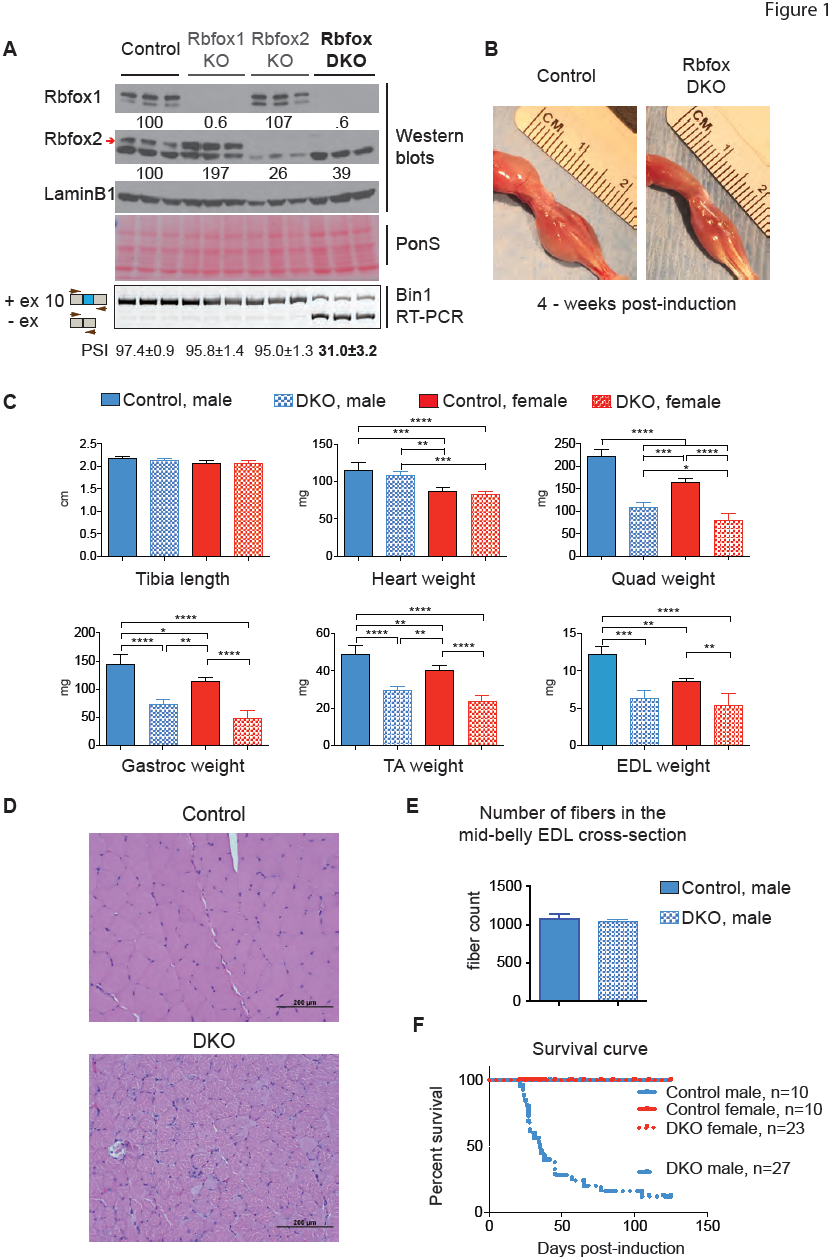
Rbfox DKO in adult skeletal muscle causes rapid loss of muscle mass by reduction in fiber size. A). Western blots for Rbfox1, Rbfox2, and LaminB1 from control (Rbfox1^f/f^, Rbfox2^f/f^), Rbfox1 (Rbfox1^f/f^: ACTA1-rtTA,tetO-cre), Rbfox2 (Rbfox2^f/f^: ACTA1-rtTA,tetO-cre), and double Rbfox knockout (Rbfox1^f/f^, Rbfox2^f/f^,ACTA1-rtTA,tetO-cre) Tibialis anterior (TA) muscles. Red arrow indicates confirmed Rbfox2 band. Ponceau S staining of representative blots as additional loading control. RT-PCR using primers flanking Bin1 exon 10 (bottom panel) from the same TA muscle that were used for westerns. Percent Spliced In (PSI) values are indicated below (mean ± Standard Deviation, SD from n=3). B) Representative image of the hind limbs from control and Rbfox DKO four weeks after starting the dox diet. C) Average measurements of tibia length and weights of isolated heart and different skeletal muscles: quadriceps (Quad), gastrocnemius (Gastroc), TA, and extensor digitorum longus (EDL) from control and DKO male and female mice (n ≥ 3). The columns and error bars in graphs indicate mean ± SD. p values as indicated by asterisk were calculated by ANOVA, *p ≤ 0.05, **p ≤ 0.01, ***p ≤ 0.001, and ****p ≤ 0.0001. D) Hematoxylin and eosin staining of Quadriceps from control (top) and DKO (bottom) mice 4-weeks after induction of Rbfox DKO. E) Total fiber number count from the mid-belly cross-section of EDL muscle from control and DKO mice 4-weeks after induction of knockout (n=3). F) Survival curves for control and DKO, male and female, animals.

Rbfox DKO animals displayed a marked loss of muscle mass and total body weight, compared to age-and sex-matched control animals, within four weeks of knockout induction (**Fig. 1B** and **S1A**). Tibia length was the same between aged-matched control and DKO animals (**Fig. 1C**), but there were significant reductions in the weight of five predominantly fast-twitch skeletal muscles—quadriceps, gastrocnemius, tibialis anterior (TA), extensor digitorum longus (EDL), and triceps—in both male and female mice (**Figs. 1C** and **S1A**). The weight of the soleus, which contains approximately 40% type I or slow-twitch fibers, did not decrease in mice of either sex (**Fig. S1A**), most likely because the knockout of Rbfox2 in the soleus was not as efficient as in fast-twitch muscle (data not shown). Consistent with the muscle-specificity of the Cre recombinase (Rao and Monks, 2009), we did not observe a change in heart or liver weights (**Figs. 1C** and S1A). Single knockout of Rbfox1 or Rbfox2 did not reduce skeletal muscle weight (**Fig. S1B**).

Because ACTA1-rtTA Cre recombinase expression is slightly leaky in the absence of doxycycline (dox) (Rao and Monks, 2009), we measured the weight of the gastrocnemius muscle several times between starting the dox-containing diet at the age of seven weeks until 21 days later. The DKO gastrocnemius muscle initially weighed 15% less in DKO mice than in uninduced animals, but by 21 days the DKO gastrocnemius weight was 41% less than that of control animals (**Fig. S1C**). Based on these results, we conclude that Rbfox splicing factors are required to maintain skeletal muscle mass in adult animals.

Histological analysis of Rbfox DKO quadriceps cross-sections showed non-uniform and dramatically reduced fiber size in DKO mice compared to controls (**Fig. 1D**). We observed that few myonuclei were centrally located which suggest that minimal degeneration-regeneration had occurred. We also did not observe infiltration of inflammatory cells in DKO muscles. The total number of muscle fibers in the mid-belly cross-sections of EDL muscles did not differ between DKO and control mice (**Fig. 1E**). We conclude that reduction in fiber size is the main reason for loss of muscle mass after Rbfox DKO.

Despite the overall similarity of muscle loss in the male and female mice, there was one notable difference between the sexes: 80% of the male Rbfox DKO animals died as early as three weeks after knockout induction, but none of the control or DKO female animals died during the course of our experiments (**Fig. 1F**).

### Deletion of Rbfox1 and Rbfox2 causes loss of muscle strength but not endurance

To evaluate muscle function we measured the forelimb and all-limb grip strength in DKO male and female animals 2 to 8 weeks after starting the dox-containing diet. (The premature death of the males prevented us from studying them at a later time point.) Both sexes showed significant reduction in forelimb and all-limb grip strength compared to age-and sex-matched control animals (**Fig. 2A**). We also found a significant reduction in upside-down hanging time in both male and female DKO animals. Performance on a treadmill assay revealed no significant difference in endurance between control, DKO male and DKO female mice (**Fig. 2A**). We also measured the total activity of male and female mice in the vertical (rearing) and horizontal (ambulation and fidgeting) planes. Male DKO mice displayed significantly less rearing than control mice (**Figs. S2A,B**), whereas female DKO mice displayed significantly greater horizontal plane activity and a trend for increased rearing compared to control females (**Figs. S2A,B**). The activity differences between control and DKO animals in both sexes were predominantly due to movement in the active period (12 hrs, night) (**Figs. S2A,B**).

**Figure 2.**
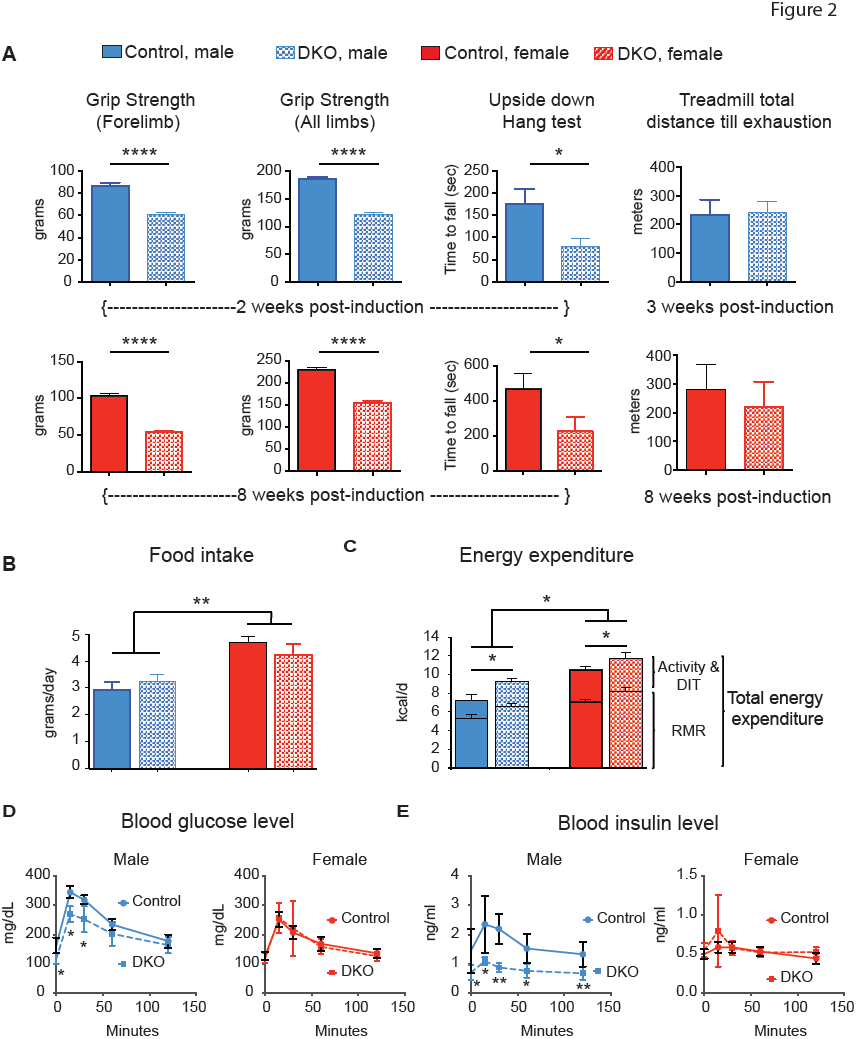
Rbfox1/2 knockout in adult muscle causes reduced muscle strength and altered glucose homeostasis. A) Forelimb (left panels) and all limb (left middle panels) grip strength in control (Rbfox1^f/f^, Rbfox2^f/f^) and Rbfox DKO (Rbfox1^f/f^, Rbfox2^f/f^,ACTA1-rtTA,tetO-cre) male (top panels) and female (bottom panels), 2-weeks and 8-weeks after knockout induction, respectively. The right middle panels show the average time animals hang upside down on wire mesh. Treadmill performance (right panels) of control and DKO mice with increasing speed till exhaustion (at 10% incline). n **>** 9 for all panels. Singly housed age-matched male and female animals were monitored for food intake (B) and indirect calorimetry was utilized to estimate total and resting energy expenditure (C) (n=5 for male, and n **>** 3 for female). Values shown are least square means ± SE adjusted using ANCOVA for differences in body weight (for food intake) and lean mass (for energy expenditure; fat mass does not contribute to variance). Abbreviation: RMR - resting metabolic rate, DIT-diet-induced thermogenesis. Activity and DIT is not significantly different between genotype or gender. (D and E) Glucose tolerance test in male (left panels) and female (right panels) mice 2 weeks after induction of Rbfox knockout. D shows blood glucose level; E shows insulin levels in control and DKO male (left) and female (right) mice (n=5). The columns and error bars in the figure indicate mean ± SD. p values were calculated by multiple Student’s t test and are indicated as follows: *p **<** 0.05, **p **<** 0.01, and ****p< 0.0001.

### Double knockout adult mice expend more energy

To understand how the loss in muscle mass affects energy metabolism, we performed indirect calorimetry in age-matched control and DKO male and female animals starting 10 days after knockout induction. We found no significant differences in 24-hr, weight-adjusted food intake between control and DKO animals, although females ate more than males (**Fig. 2B**). Both male and female DKO animals, however, expended more energy than age-matched control animals (**Fig. 2C**); this difference was attributable to differences in resting metabolic rate, and therefore was evident both in periods of rest (12 hrs, day) and activity (12hrs, night) (**Fig. S2C**).

Since skeletal muscle is a primary organ for maintaining glucose homeostasis, we performed glucose tolerance tests to determine the effect of Rbfox DKO on glucose metabolism in age-matched control and DKO animals two weeks after knockout induction. Before glucose injection, male but not female DKO mice had lower levels of serum glucose relative to control animals. Even after glucose injection, glucose levels remained lower in DKO male animals but not in female animals (**Fig. 2D**). Serum insulin levels were nearly half that of controls in the DKO male animals while DKO females did not show a difference (**Fig. 2E**). These results indicate that glucose homeostasis is altered in male Rbfox DKO animals and that male DKO animals show a trend towards insulin hypersensitivity.

### Deletion of Rbfox1 and Rbfox2 causes widespread transcriptome changes

To investigate the molecular mechanism by which Rbfox1/2 deletion produced these changes, we performed 100-bp paired-end RNA-sequencing (RNA-seq) using polyadenylated RNA from the EDL and soleus muscles from two male DKO mice, two weeks after knockout induction, and two age-matched male controls. For each sample we obtained over 170 million reads, 87% of which mapped to the genome (Table 1 in supplementary information). Mapped reads were used to quantify transcriptome changes including gene expression and alternative use of first, last, and internal exons. Gene expression changes for biological replicates correlated very highly (R^2^ = 0.99) demonstrating the reproducibility of RNA-seq and computational analysis of the data (**Fig. S3A**). In the DKO EDL, which showed significant loss of muscle mass, 832 genes were differentially expressed (FDR <0.05) with 495 genes (~60%) showing upregulation and 337 genes (~40%) showing downregulation when compared to control animals (**Fig. 3A** **and Table S1**). In the DKO soleus, which did not show significant changes in mass in males or females, 206 genes were differently expressed (FDR <0.05), with 132 genes showing upregulation and 74 genes showing downregulation relative to their levels in control animals (**Fig. 3A** **and Table S1**).

**Figure 3.**
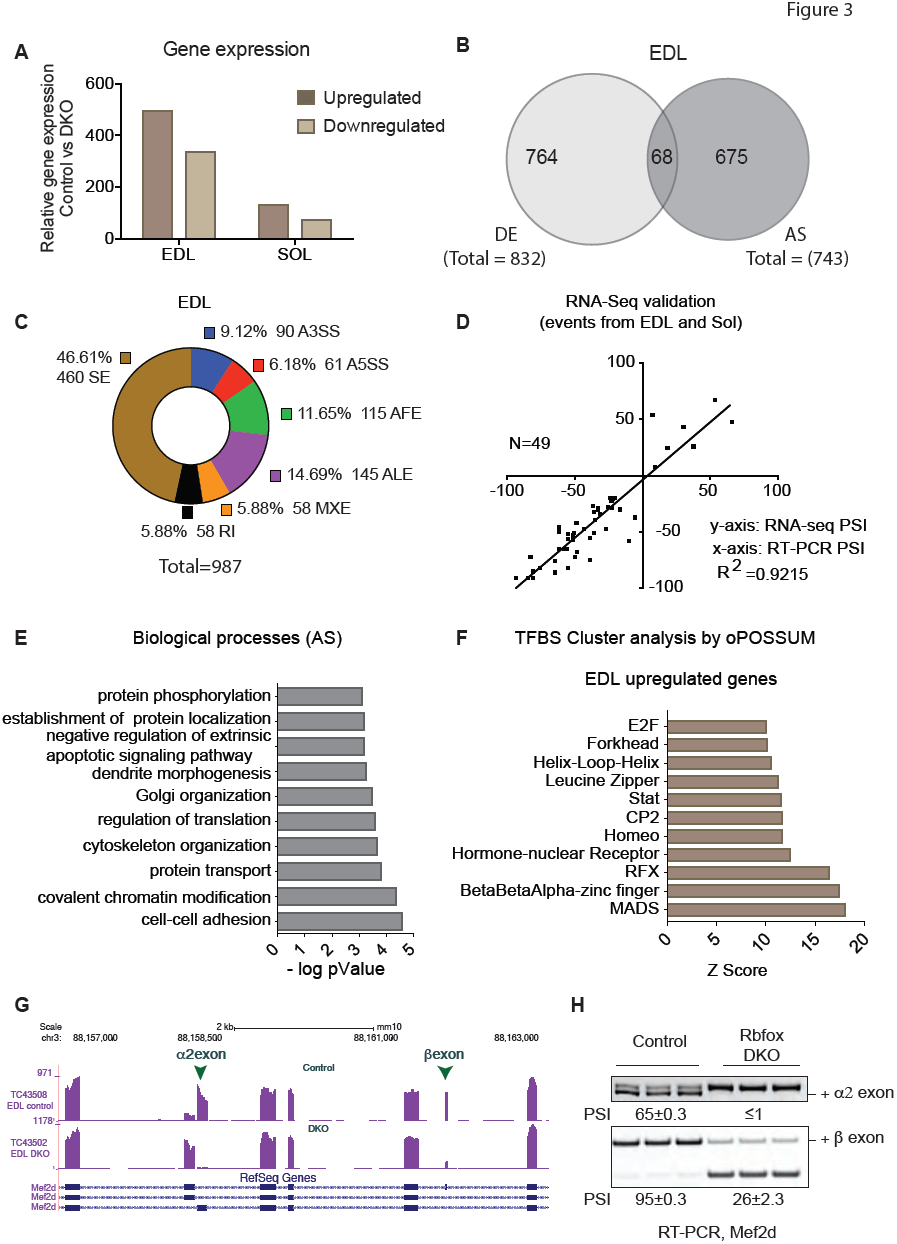
Rbfox knockout in skeletal muscle causes wide-spread transcriptome changes. A) Bar graphs showing gene expression changes (FDR <0.05) in EDL and soleus, two weeks after induction of Rbfox DKO compared to control muscles. B) Venn diagram showing differentially expressed (DE) and alternatively spliced (AS) gene transcripts in DKO in comparison to control EDL muscle (FDR <0.05 for DE and PSI≥ 0.2 for AS). C) Donut chart showing description of transcriptome changes in EDL muscle, 2 weeks after Rbfox knockout, compared to control muscle. Abbreviations: SE - cassette exons, A3SS - alternative 3’ splice site, A5SS - alternative 5’ splice site, AFE - alternative first exon, ALE - alternative last exon, MXE - mutually exclusive exons, RI - retained intron. D) Forty-nine randomly selected alternative splicing events in EDL and soleus muscles were validated by RT-PCR and APSI was plotted against APSI derived from RNA-seq data. E) DAVID gene ontology (GO) analyses showing biological processes that are enriched in genes those are alternatively spliced in Rbfox knockout muscles. F) oPOSSUM analysis to determine the enrichment of clustered transcription factor binding sites (TFBS) in genes that are upregulated after Rbfox knockout. G) RNA-seq tracks for Mef2d in control and Rbfox DKO muscles showing that muscle-specific inclusion α2 and β exon requires Rbfox proteins. H) RT-PCR showing reduced PSI for Mef2d α2 and βexons in DKO muscle when compared to control muscle (n=3).

When we examined alternative splicing, we found that 743 genes showed different usage of exons between control and Rbfox DKO EDL muscle with a Percent Spliced In (PSI) cutoff of ≥20% (**Fig. 3B** **and Table S2**)(Wang et al., 2008). There is little overlap, of only 68 genes, between genes that change in overall mRNA levels (832 genes) and those that show differential usage of alternative exons (743 genes) (**Fig. 3B**). Similarly, of the 389 genes that show alternative use of exons, only 9 differed in overall expression (FDR <0.05) between DKO and control soleus muscle (**Fig. S3B** **and Table S2**). The majority of alternative exons altered by Rbfox DKO are cassette-type in both EDL (460 exons) and soleus (146 exons) (**Figs. 3C** and **S3C**). RT-PCR analysis of splicing for 49 alternative exons found a strong correlation (R^2^ = 0.92) between ΔPSI from RT-PCR and computed from the RNA-seq data (**Fig. 3D**).

We used the DAVID functional annotation tool to identify biological processes that are enriched among these groups of genes (Huang da et al., 2009). Functions that are enriched among those genes that are downregulated include oxidation-reduction, calcium handling, and cell signaling (**Fig. S3D**). Enriched functions among upregulated genes include muscle differentiation and development, muscle contraction, cell adhesion, and transcription regulation (**Fig. S3E**). Enriched categories among genes showing altered splicing include cell-cell adhesion, cytoskeleton organization, chromatin modification, and regulation of protein metabolism including translation, transport, localization, and phosphorylation (**Fig. 3E**).

We used a web-based tool, oPOSSUM-3, to identify over-represented clustering of Transcription Factor Binding Sites (TFBS) within 5000 bases upstream and downstream of genes that show altered expression in Rbfox DKO muscle (Kwon et al., 2012). In upregulated genes, TFBS clustering analysis showed enrichment for the MADS family of transcription factors (**Fig. 3F**). TFBS analysis identified 441 MADS binding sites clustered in the promoter regions of 206 genes that are upregulated, but only 19 binding sites in 17 genes that are downregulated in Rbfox DKO muscle; this suggests that alterations in the MADS transcriptional program leads to upregulation of its targets (**Table S3**).

It is worth noting that the MADS family includes the Mef2 transcription factors. We and others have shown that Mef2a and Mef2d splicing is coordinately and directly regulated by Rbfox proteins during myogenesis in culture (Runfola et al., 2015; Singh et al., 2014); we have also previously shown that the Mef2d splice variant expressed in adult skeletal muscle is required for late stages of muscle differentiation (Singh et al., 2014). Here we found that Rbfox DKO muscle displayed significantly lower use of the β exon of Mef2a and both the α2 and β exons of Mef2d (**Figs. 3G-H** and **S3G-H**), representing a loss of muscle-specific isoforms. We conclude that Rbfox coordinately regulates splicing of Mef2a and Mef2d to maintain the expression and proper activity of muscle-specific isoforms (Sebastian et al., 2013).

### Altered proteostasis in Rbfox DKO muscle

Skeletal muscle mass is determined by the balance between rates of protein synthesis and protein degradation. The rate of protein synthesis in skeletal muscle is regulated by growth factor, amino acid, and insulin signaling pathways (**Fig. 4A**). We measured the rate of protein synthesis in control and Rbfox DKO TA muscles two weeks after knockout induction using puromycin in the SUnSET assay (Goodman and Hornberger, 2013; Schmidt et al., 2009). There was no reduction of puromycin-labeled peptides in DKO muscle compared to controls (**Fig. 4B**), indicating that the loss of muscle in Rbfox DKO skeletal muscle was not due to a reduction in protein synthesis. We therefore turned our attention to proteolytic mechanisms of muscle loss.

**Figure 4.**
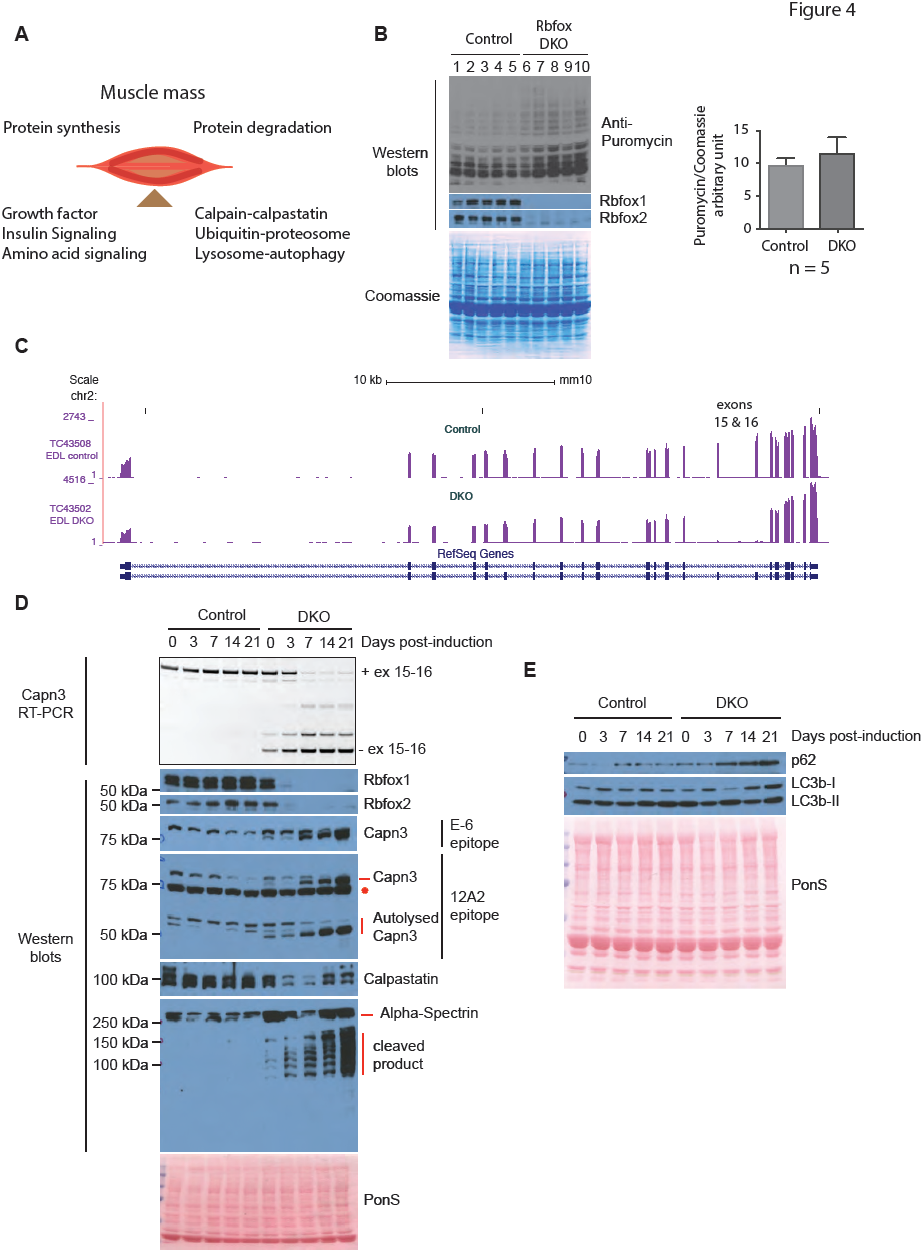
Altered regulation of Calpain in Rbfox DKO muscles. A) Maintenance of skeletal muscle mass is a balance between protein synthesis and protein degradation. Major determinants of protein synthesis and protein degradation are also indicated. B) Puromycin incorporation into nascent proteins in TA muscles was quantified in five control (lanes 1–5) and Rbfox DKO animals (lanes 6–10) 2-weeks after induction of Rbfox DKO. Western blots using antibody against puromycin or Rbfox1 or Rbfox2, and Coomassie staining of the blot are shown. Puromycin intensity/total Commassie signal was calculated as a relative measure of total protein synthesis, and plotted in a bar graph (right panel). The columns and error bars in graphs indicate mean ± SD. C) RNA-seq tracks showing Capn3 in control and Rbfox DKO muscles. Exon 15 and 16 is nearly completely skipped in Rbfox DKO muscles. D) RT-PCR for Calpain 3 using primer pairs annealing in exons 14 and 17, and Western blots using antibodies against indicated proteins in uninduced (0 day) and 3,7,14, and 21 days induced TA muscles from control and DKO animals. The same TA muscles were used to prepare RNA and protein extracts for RT-PCR and Western blots. E) Western blots for indicated proteins to assess the effect of Rbfox DKO on autophagy.

Protein degradation is tightly regulated by three major proteolytic systems: 1) Calpain-Calpastatin, 2) lysosome-autophagy, and 3) ubiquitin-proteasome (**Fig. 4A**). We focused our analysis on alternatively spliced genes involved in the three major protein degradation systems. We prioritized targets based on the extent of splicing changes in DKO based on RNA-seq data and their binding to Rbfox in muscle culture based on iCLIP data (Singh et al., 2014). We found that two consecutive alternative exons (exons 15 and 16) in Capn 3 transcripts were almost completely skipped in Rbfox DKO muscle (**Fig. 4C**). Autolytic cleavage within the IS1 region enables proteolytic Capn3 activity, and cleavage within the IS2 region of Capn3 is required for complete degradation and inactivation of Capn3 (**Fig. S4A**) (Ono et al., 2014). Skipping of exons 15 and 16 in Rbfox DKO muscle likely produces a Capn3 isoform that lacks the cleavage site within the IS2 region, thereby stabilizing active Capn3 causing it to accumulate. We hypothesized that this contributes to severe muscle loss by tilting the protein balance towards protein degradation.

To determine whether the timing of Rbfox loss correlated with altered splicing of Capn3 and initiation of muscle loss, we performed western blots for Rbfox1 and Rbfox2 and RT-PCR for splicing of Capn3 exons 15 and 16 using primers complementary to exons 13 and 17 in uninduced and DKO muscles at 3, 7, 14, and 21 days after starting the dox diet. Rbfox1 and Rbfox2 knockout was efficient 3 days after starting the dox diet, which correlates with increasing expression of Capn3 transcripts lacking exons 15 and 16 (**Fig. 4D**). By 7 days post-induction, the majority of the mRNA expressed from the *Capn3* gene lacks both exons. We performed western blots for Capn3 using two antibodies, one recognizing an epitope in the first 80 amino acids (E-6, Santa Cruz) and the other (12A2, Leica) recognizing an epitope within residues 355–370 (**Fig. S4A**). Both antibodies recognize the full-length protein and 12A2 also detects the enzymatically active (autolysed) C-terminal fragment of Capn3 (Charton et al., 2016; Fanin et al., 2003; Richard et al., 1995). Both antibodies recognized a band corresponding to full-length protein at 94 kDa in control animals and a slightly smaller band predominating following Rbfox DKO, likely encoded by the mRNA lacking exons 15 and 16 (**Fig. 4D**). Importantly, both the full-length isoform and the active autolysed C-terminal fragments (55 and 60 kDa) detected by 12A2 accumulated after induction of the DKO, correlating with the timing of rapid muscle loss. These results indicate that loss of Rbfox proteins caused altered splicing and production of smaller isoforms of Capn3, which is autolytically cleaved, proteolytically active, and stably expressed in DKO muscles.

We next determined whether Capn3 substrates are degraded in Rbfox DKO muscle. Several studies have used calpastatin as a Capn3 substrate (Ono et al., 2004; Ono et al., 2014). Calpastatin is also an endogenous inhibitor of Capn1 and Capn2; we predict that reduced calpastatin levels in Rbfox DKO would lead to increased Capn1 and Capn2 activity. By western blotting, we found that calpastatin is stably expressed in control muscles, but it was reduced in DKO muscles within 3 days after starting the dox diet (**Fig. 4D**). Alpha-spectrin is another substrate for Capn3 (and possibly Capn1 or Capn2) (Huang and Forsberg, 1998; Nakajima et al., 2006; Takamure et al., 2005; Zhang and Bhavnani, 2006), and western blotting showed a prominent band at the expected size for alpha-spectrin in control muscles, but an increasing proportion of several smaller fragments appeared in DKO muscles after starting mice on the dox diet (**Fig. 4D**). These results support the hypothesis that increased calpain activity in Rbfox DKO muscle causes degradation of calpastatin and alpha-spectrin, and likely other calpain substrates, which in turn contributes to loss of muscle mass.

Several cases of muscle atrophy have been associated with upregulation of E3 ubiquitin ligases such Fbxo32 (atrogin-1) and Trim63 (MuRF1), which target sarcomeric components for degradation (Bodine and Baehr, 2014; Bodine et al., 2001). We performed western blotting for Murf-1, Trim 55 (MuRF-2), Trim 54 (MuRF-3), and Atrogin-1, and found that Murf-1, Murf-3, and Atrogin-1 are expressed at similar levels in control mice and DKO mice after induction (**Fig. S4B**). Murf-2 levels decreased slightly in the DKO but overall we did not observe any increase in E3 ubiquitin ligases that are known to be upregulated in several muscle atrophy conditions.

To assess the contribution of autophagy to muscle loss in Rbfox DKO muscle, we performed western blotting of several proteins in control and DKO animals at different time-points after DKO induction. In lysosome-autophagy proteolytic pathways, the autophagosome must fuse with the lysosome for substrate degradation to occur. Autophagosome formation requires the activity of the microtubule-associated protein 1A/1B-light chain 3 beta (LC3B), which is first proteolytically cleaved to form LC3B-1. LC3B-I is then converted to LC3B-II by lipidation, which is present in the autophagosome membrane. Thus, different LC3B isoforms are used as markers of autophagic activity (Mizushima et al., 2010). We also quantified levels of Sequestrome1 (Sqstm1 or p62), a ubiquitin-binding protein and adaptor molecule that binds and sequesters substrates into autophagosomes (Mizushima et al., 2010). Compared to control muscle, we found a trend for accumulation of p62 and LC3B in DKO muscle over time (**Fig. 4E**).

To characterize the lysosome-autophagy flux *in vivo* in Rbfox DKO muscles, we injected mice with colchicine, a drug that depolymerizes microtubules and impairs autophagy by inhibiting autophagosome-lysosome fusion (Ju et al., 2010). Mice were injected with colchicine or vehicle (water) for two consecutive days at day 4–5(early) or day 12–13(late) time-points after DKO induction and the tibialis anterior was harvested 48 hours after first injection, at day 6 and day 14, respectively. At the early time-point, there was no difference in the levels of p62 or LC3B-II between water-injected control and Rbfox DKO mice. Compared to water-injected mice, colchicine-injected mice without Rbfox KO showed the predicted accumulation of LC3B-II and p62 at early time-point (**Fig. 5 A-B**). In contrast, colchicine injected Rbfox DKO mice failed to produce an increase in p62 or LC3B-II (**Fig. 5 A-B**), indicating an early Rbfox-dependent reduction in autophagy flux.

**Figure 5.**
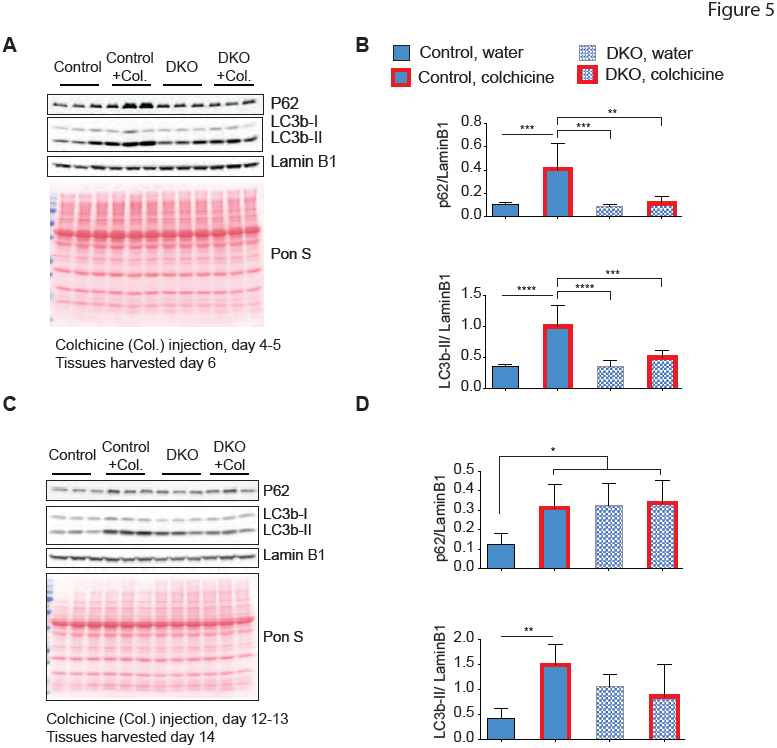
Autophagy flux is reduced in Rbfox DKO muscle. A) Western blots using antibodies against indicated proteins from TA muscles from control and DKO muscles. Control (Con) or Rbfox knockout (DKO) mice were injected with Colchicine (Col) or water at day 4–5 (A) or day 12–13 (C), and muscles were harvested the next day for western blotting. Levels p62 and LC3b-II levels were normalized to Lamin B1 and plotted in B and D, for western blots in A and C, respectively (n=4 for early time-point and n > 3 for late time-point). The columns and error bars in graphs indicate normalized mean intensity of p62 or LC3b-II ± SD. Red lines in the graph columns correspond to values from Col injected mice. p values as indicated by asterisk were calculated by ANOVA, *p ≤ 0.05, **p ≤ 0.01, ***p ≤ 0.001, and ****p ≤ 0.0001.

This early reduction in autophagy flux is expected to increase the accumulation of p62 and LC3B-II over time, and consistently, we observed substantially increased p62 and LC3B-II levels in water-injected DKO mice in comparison to control mice, two weeks after induction of KO. Compared to water-injected mice, colchicine-injected mice without Rbfox KO showed the predicted accumulation of LC3B-II and p62 at late time-point (**Fig. 5 C-D**). Whereas in DKO mice, already high levels of LC3B-II and p62 did not increase further after colchicine injection (**Fig. 5 C-D**), confirming that early Rbfox-dependent reduction in autophagy flux is maintained at later time-point. These results indicate that there is a marked reduction in autophagy flux in Rbfox DKO as early as day 4–6 after KO induction and which persists at later times.

From these experiments, we conclude that knockout of Rbfox proteins in skeletal muscle leads to increased calpain activity and impaired autophagic flux, thereby altering two key proteolytic systems in skeletal muscle. Such a dramatic imbalance of protein synthesis and degradation will result in the net loss of protein and promote atrophy.

## DISCUSSION

We have found that concurrent deletion of Rbfox1 and Rbfox2 in adult mouse skeletal muscle causes the dramatic and rapid loss of muscle mass. This response directly correlated with impaired autophagy flux and the formation of alternative isoforms of calpain3 that are not degraded rapidly, leading to the accumulation of calpain3. Within four weeks of inducing the DKO, both male and female mice lost nearly half the muscle mass in the five fast-twitch muscles we measured. We also observed a significant decrease in whole-body lean mass (data not shown) after DKO induction indicating ubiquitous loss of muscle mass in both male and female Rbfox DKO mice. Despite the comparable muscle loss, however, only male mice had a shortened lifespan. The cause of this premature lethality is unclear, but we found alterations in glucose metabolism, trending towards insulin hypersensitivity, only in male animals. Interestingly, deletion of the estrogen receptor alpha (ERα) specifically in skeletal muscle of female mice causes alteration in glucose homeostasis, increased fat accumulation, and impaired mitochondrial function (Ribas et al., 2016), so it may be that estrogen protects the female mice from death despite dramatic loss of muscle mass after induction of Rbfox knockout.

Inhibition of lysosome-autophagy causes loss of muscle mass in humans and mice, emphasizing the importance of autophagy in muscle growth, remodeling, and maintenance. Loss-of-function mutations in lysosome-associated membrane protein 2 (LAMP-2), a protein important for fusion of autophagosome with lysosome, underlie Danon disease, in which patients present with skeletal muscle weakness, intellectual disability, and accumulation of vacuoles with autophagic material in the heart and skeletal muscle (Bell et al., 2016a; Masiero and Sandri, 2010; Raben et al., 2008; Tanaka et al., 2000). Deletion of Autophagy-related protein 7 (ATG7), a protein required for autophagosome formation, in skeletal muscle causes loss of muscle mass in mice (Masiero et al., 2009). These mice show increased expression of atrogin-1 and Murf1, and loss of muscle mass, which is caused in part by compensatory upregulation of ubiquitin-proteasome proteolytic activity. We found that in Rbfox DKO caused a rapid reduction in autophagy flux and led to greater accumulation of p62 and LC3b-II in DKO muscles over time (**Fig 5A-D**). The reduction in autophagy flux in combination with increased calpain activity is expected to increase the demand for proteolytic activity. We propose this compensatory increase in proteolytic activity also contributes to the loss of muscle mass in Rbfox DKO animals.

An isoform of Capn3 lacking IS1 or IS2 is proposed to be dominantly active with catastrophic effects (Beckmann and Spencer, 2008; Spencer et al., 2002). Consistent with this notion, muscle-specific overexpression of full-length Capn3 at high (over 25x wild-type) levels yields no observable phenotype, but overexpression of Capn3 lacking exon 6, which encodes most of the IS1 region, produces a severe phenotype that includes reduced muscle mass, smaller muscle fibers, kyphosis, and early death. The authors concluded that the phenotype is due to altered activity of Capn3 during the postnatal development (Kramerova et al., 2004; Spencer et al., 2002). Overexpression of a Capn3 cDNA lacking exon 15, which only lacks 6 amino acids in the IS2 region, leads to a subtle phenotype in the soleus, apparently without affecting other muscles, and Capn3 protease activity towards casein was much higher for muscle expressing the isoform lacking exon 15 compared to wild-type Capn3 or a Capn3 isoform lacking the full IS1 domain due to missing exon 6 (Spencer et al., 2002). In our study, we induced Rbfox DKO in animals older than 7 weeks so that we would not affect the normal postnatal function of Capn3. Rbfox knockout switched the expression of full-length Capn3 to an isoform lacking both exon 15 and 16 (114 nucleotides) that encodes the majority (44 of 71 amino acids) of the IS2 region, which is required for complete degradation of Capn3 (**Fig. S4**). The smaller Capn3 isoform is autolytically processed to generate a catalytically active form of Capn3, and we propose that stable expression of the shorter, active form of Capn3 contributes to muscle loss in Rbfox DKO mice.

Calpains are nonprocessive proteases that cleave substrates at specific sites (Ono and Sorimachi, 2012). It has been predicted that over 1000 proteins are calpain targets, yet fewer than 100 substrates have mapped calpain cleavage sites because calpain cleaved fragments are shortlived (Ono and Sorimachi, 2012; Piatkov et al., 2014). Despite the critical role Capn3 plays in skeletal muscle biology, it is not clear which proteins are *in vivo* substrates for Capn3: the majority of its substrates have been identified through biochemical or cell culture studies (Huang and Forsberg, 1998; Nakajima et al., 2006; Ono et al., 2004; Ono et al., 2014; Takamure et al., 2005; Taveau et al., 2003; Zhang and Bhavnani, 2006). We found that levels of two of the previously identified substrates, calpastatin and alpha-spectrin, are reduced within 7 days after induction of Rbfox knockout. Reduced calpastatin protein level would imply less inhibition of Capn1 and 2, strongly supporting a hypothesis that Capn3 is the master regulator of the calpain-calpastatin system in skeletal muscle (Ono et al., 2004). It is interesting that increased levels of Calpn3 lacking exons 15 and 16 are as detrimental to myofibers as loss of function mutations in Capn3, which are the most common cause of Limb Girdle Muscular Dystrophy Type 2A (LGMD2A), a childhood-onset, progressive disease. The dramatic skeletal muscle phenotype in the Rbfox DKO mouse provides clear evidence not only of the importance of alternative splicing for maintenance of adult muscle mass, but also of the sensitivity of muscle to alterations in proteostasis as has been observed for the brain (Navone et al., 2015; Smith et al., 2015; Sulistio and Heese, 2016).

## STAR Methods

### Mice

The Institutional Animal Care and Use Committee at Baylor College of Medicine approved all animal procedures. All mouse lines in this study were obtained from Jackson Laboratories and in C57BL/6J genetic background (Bar Harbor, ME). Previously described floxed Rbfox lines were mated with a muscle-specific, doxycycline inducible *Cre* line to generate single Rbfox1 (Rbfox1^f/f^: *ACTA1-rtTA^cre/+^*) or Rbfox2 (Rbfox2^f/f^: *ACTA1-rtTA^cre/+^*) and double Rbfox (Rbfox1^f/f^: Rbfox2^f/f^: *ACTA1-rtTA^cre/+^*) knockout (Gehman et al., 2012; Gehman et al., 2011; Pedrotti et al., 2015; Rao and Monks, 2009). For inducing the knockout, 7-weeks or older control (floxed mice) and littermate single and/or double knockout mice were fed on doxycycline-containing diet (2 gm/kg, Bio-Serv, NJ) for one week and then transferred to normal chow (5V5R, LabDiet, MO). Mice were anesthetized with isoflurane and tissues were isolated, weighed, and snap frozen in liquid nitrogen for further analysis.

### Mouse muscle strength and activity assays

Treadmill assays were performed on the Exer 3/6 six treadmill (Columbus Instruments, OH) at a starting speed of 6 meters/min for first two minutes (10 degree incline) increased 2 meters/min every two minutes until exhaustion. The speed and time were used to measure the total distance run by mice in each group. Grip strength was performed using a 1027SM grip strength meter with single sensor (Columbus Instruments, OH). Forelimb and all limb grip strength used 5 measurements of grams of force produced in the t-pk mode and average values were calculated. For the hang test, soft padding was added in cardboard box (14 in x14 in x14 in), a wire grid was placed on top of the box and the mouse was placed upside down; the time until the mouse could no longer hold onto the wire was recorded for each mouse.

### Histological analyses

For histology of muscles, skin and facia was removed and whole limbs were fixed overnight in 10% buffered formalin. Muscles were isolated and processed for paraffin embedding, routine sectioning, hematoxylin and eosin staining.

### Food Intake, Energy Expenditure, Activity

The Comprehensive Laboratory Animal Monitoring System (Columbus Instruments, OH) was used to measure food intake, activity, and energy expenditure. Mice were adapted to the cages at 9 days after starting the dox diet and various measurements were taken from day 13 to day 17 to quantify energy balance, food intake, and activity as previously described (Crossland et al., 2017).

### Glucose Tolerance Test

The glucose tolerance test was performed by the Mouse Metabolism Core at Baylor College of Medicine. Two weeks after starting the dox diet, mice were starved for 6 hours, and then glucose (1.5 g/kg of body weight) was injected intraperitoneally. Blood was drawn at the indicated times after injection and glucose and insulin levels were measured as described before (Saha et al., 2010).

### RNA isolation, RNA-seq, and Analysis of RNA-seq data

RNA was isolated from mouse muscles using TRIzol reagent (Invitrogen) or RNeasy fibrous tissue mini-kit (Qiagen, Germantown, MD). We used RNA from two biological replicates from control and Rbfox DKO EDL and soleus muscle at 14 days after knockout induction. The poly(A) RNA were isolated for paired-end 100bp sequencing as previously described (Singh et al., 2014). The analysis of the RNA-seq data was performed by AccuraScience LLC (Johnston, IA). RNA-seq reads were aligned to the genome using TopHat, Edge R was used for quantification of gene expression changes, MISO was used for quantification of alternative splicing changes, and Percent Spliced In (PSI) was calculated, as previously described (Katz et al., 2010; Robinson et al., 2010; Trapnell et al., 2010; Wang et al., 2008). To validate RNA-seq alternative splicing data, we performed standard RT-PCR using primers that flank specific alternative exons and PCR products were separated on a 5% acrylamide gel. The gel was imaged and PSI was calculated as previously described (Singh et al., 2014).

### Western blotting

Frozen muscles were homogenized with Bullet Blender 24 (Next Advance, Averill Park, NY) using zirconium oxide beads in buffer containing 50 mM Tris-HCL, pH 7.5, 100mM NaCl, 10 mM EDTA, 10 mM EGTA, 10% glycerol, 1% NP40, 50 mM NaF, 10uM MG132, 1mM PMSF, 0.5% Sodium deoxycholate, 1% SDS, protease and phosphatase inhibitor (Roche Diagnostics, Indianapolis, IN). Relative protein amount was quantified using the Pierce BCA Protein Assay Kit and equivalent amount of proteins were loaded on SDS-PAGE gels. Separated proteins were transferred to PVDF membrane. Membranes were stained with Ponceau S to assess even transfer and subsequently probed with specific primary antibodies. Secondary antibodies were conjugated to HRP and blots were imaged with ChemiDoc XRS+ (Bio-Rad, Hercules, CA) or standard autoradiography film. For quantification of western blot, images from ChemiDoc were exported for analysis and analyzed by Carestream software (Rochester, NY).

### Detection of puromycin-labeled peptides

Rbfox knockout was induced in 7-week-old control and Rbfox DKO male animals, and 2-weekslater, mice were injected intraperitoneally (IP) with Puromycin (0.04 μmol/g body weight) or water (Goodman and Hornberger, 2013; Schmidt et al., 2009). TA muscle was harvested 45 minutes after injection for western blotting with anti-Puromycin [3RH11] antibody (Kerafast, MA). The same Western blots were stained with the Coomassie for normalization of Puromycin signal by total protein signal by densitometry.

### Autophagy flux experiment

Autophagy flux was determined as described (Ju et al., 2010). Mice were injected IP with 0.4 mg/kg of colchicine (Sigma, St. Louis, MO) for two consecutive days at 24-hour interval and sacrificed approximately 48 hours after the first injection. The relative levels of LC3B-II and p62 were quantified on western blots and normalized to Lamin B1.

## ACCESSION NUMBERS

RNA-seq data is available for download from NCBI Gene Expression Omnibus (http://www.ncbi.nlm.nih.gov/geo/) under accession number xxxxxxx.

## ACKNOWLEDGEMENTS

This project was supported by the Mouse Phenotyping Core at Baylor College of Medicine with funding from the NIH (UM1HG006348); measurements of body composition, energy balance, and food intake were performed in the Mouse Metabolic Research Unit at the USDA/ARS Children’s Nutrition Research Center, Baylor College of Medicine (www.bcm.edu/cnrc/mmru) and supported by funds from the USDA ARS (USDA CRIS 6250-51000-054). RKS is supported by Post-doctoral fellowship and Scientist Development Grant from the American Heart Association (12POST11770017 and 15SDG25610021). This project is funded by grants from the Muscular Dystrophy Association and National Institutes of Health to TAC (R01HL045565, R01AR060733, and R01AR045653). We thank A. Vainshtein and V. Brandt for helpful suggestions and careful reading of the manuscript. We also thank members of the Cooper lab for discussion and help throughout the project.

## Authors’ contributions

RKS and TAC conceived the study, designed the experiments, interpreted data, and wrote the manuscript. RKS also performed the experiments. AMK helped with mouse muscle function tests and validation of RNA-seq data, and MLF contributed to the design, data analysis, and interpretation of energy balance studies.

---

**Figure S1.**
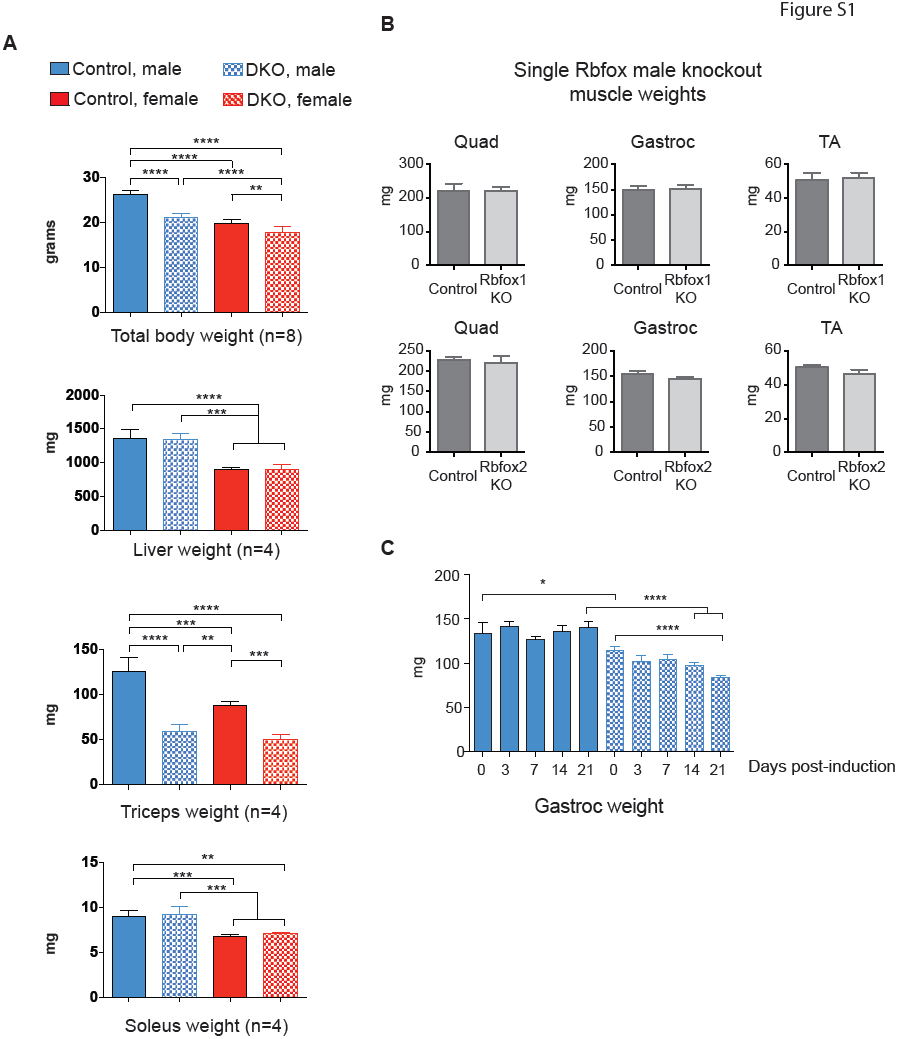
Deletion of Rbfox1 and Rbfox2 in adult skeletal muscle causes loss of muscle mass. A) Measurements of total body weight, and isolated liver, triceps and soleus muscle weights from control and DKO male and female mice (4-weeks after induction of Rbfox knockout). B) Isolated muscle weights from control and single Rbfox1 (top) or Rbfox2 (bottom) knockout male mice. C) Isolated gastrocnemius muscle weight from uninduced and 3, 7, 14, and 21 days after starting doxycycline diet. N is ≥ 3 for all experiments in this figure. The columns and error bars in graphs indicate mean ± SD. p values as indicated by asterisk were calculated by ANOVA, *p ≤ 0.05, **p ≤ 0.01, ***p ≤ 0.001, and ****p ≤ 0.0001.

**Figure S2.**
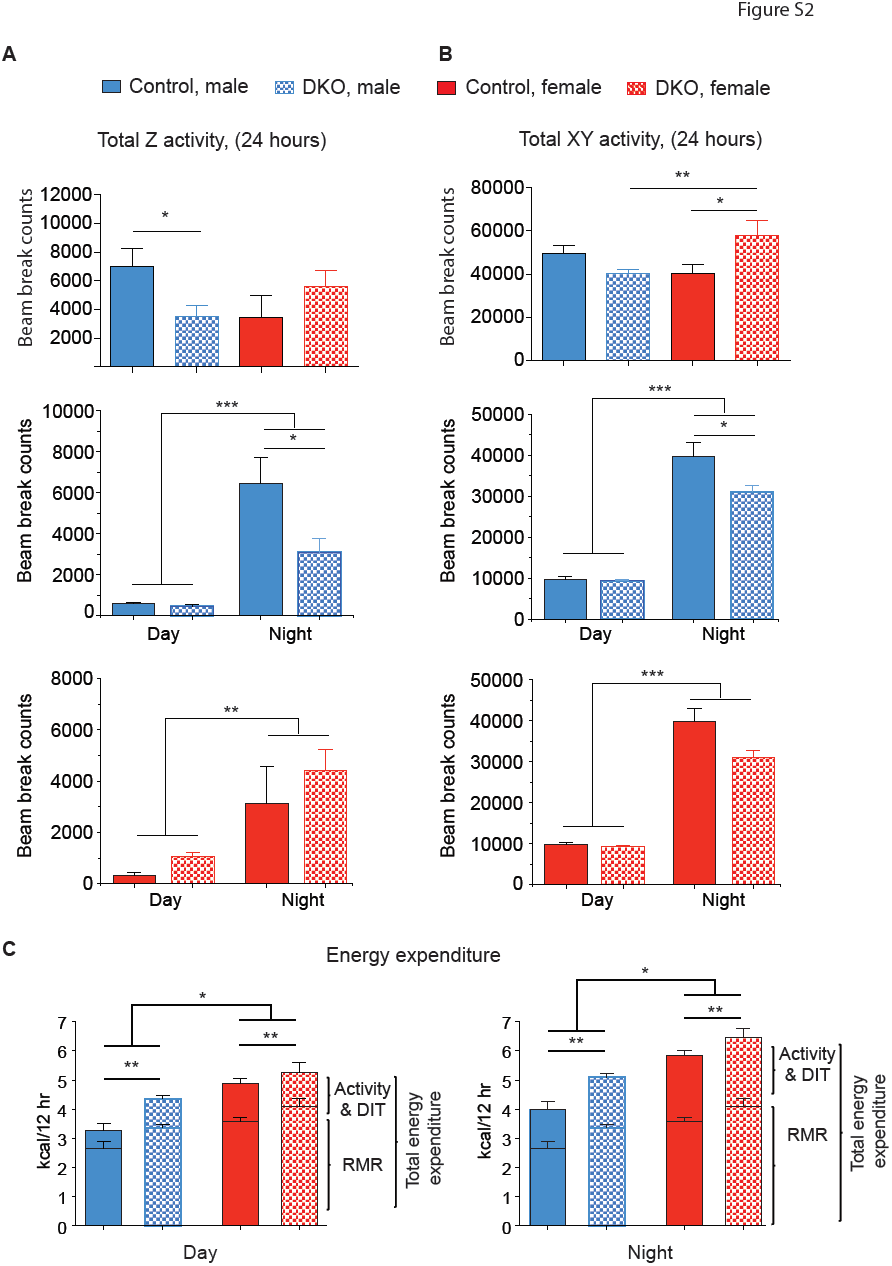
Activity and energy expenditure of control and Rbfox DKO male and female mice approximately 2 weeks after knockout induction. A-B) Average 24-hour rearing or Z-plane activity (A) and horizontal-plane activity (B) in male and female control and Rbfox DKO animals (n=5 for male, n≥3 for female). Activity data in A and B was divided into 12-hour day (rest period, 6AM to 6PM) and night (active period, 6PM to 6AM) for male (middle panels) and female (bottom panels). For the analysis of effects of time of day; we used a repeated measure ANOVA with time of day as the repeated measure for each mouse. C) Energy expenditure (adjusted for lean mass) from control and DKO mice for day (left panel) and night (right panel) are indicated (n=5 for male, n≥3 for female). Abbreviation: RMR - resting metabolic rate, DIT-diet induced thermogenesis. Activity and DIT is not significantly different between genotype or gender. The columns and error bars in graphs indicate means ± SE. p values as indicated by Asterisk were calculated by ANOVA, *p ≤ 0.05, **p ≤ 0.01, ***p ≤ 0.001.

**Figure S3.**
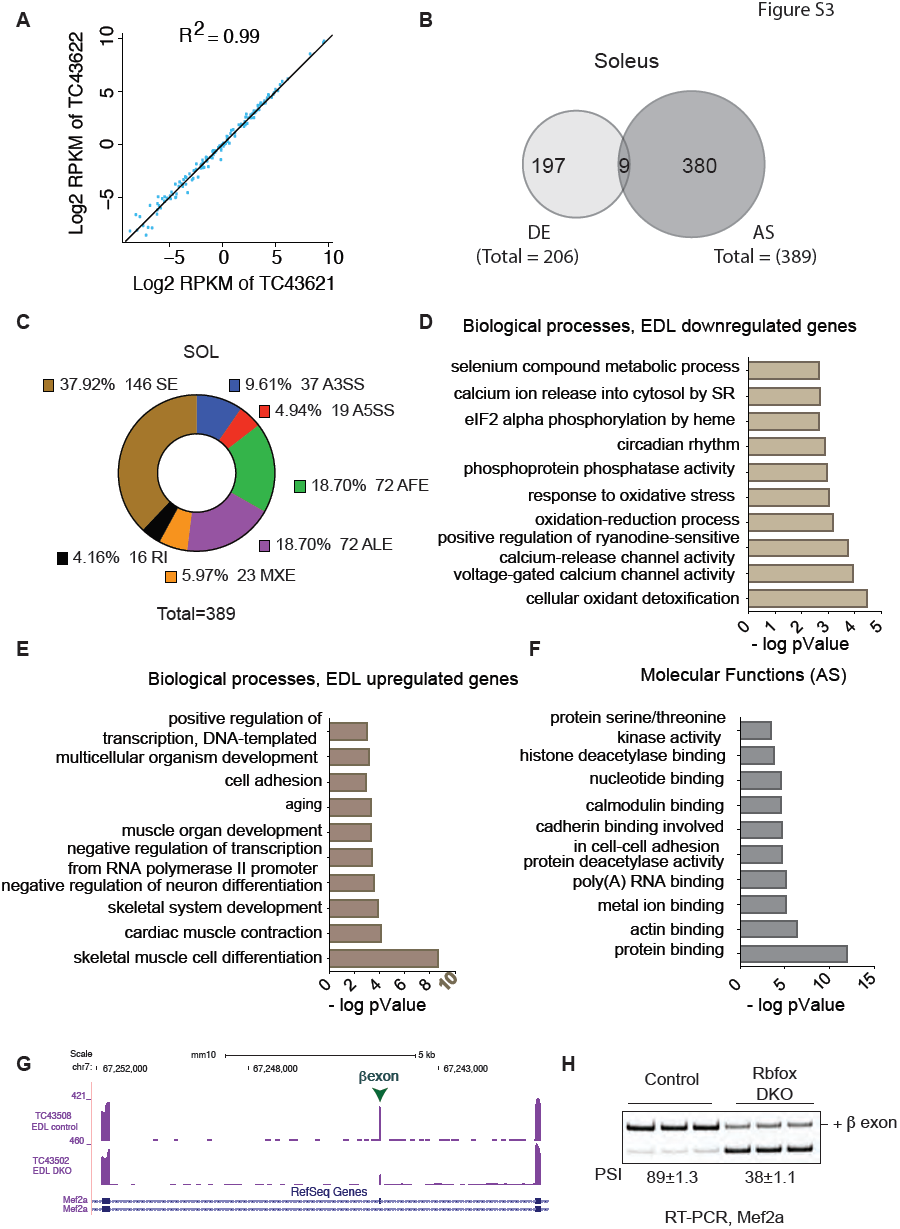
Rbfox knockout in skeletal muscle causes wide-spread transcriptome changes. A) Gene expression from two biological replicates was compared with each other. B) Venn diagram showing differential gene expression (DE) and alternative splicing (AS) changes in Rbfox knockout soleus muscles when compared to control animals at 2 weeks after knockout induction. C) Donut chart showing description of transcriptome changes in soleus muscle. Abbreviation: SE-Cassette exons, A3SS-alternative 3’ splice site, A5SS - alternative 5’ splice site, AFE-alternative first exon, ALE-alternative last exon, MXE - mutually exclusive exons, RI - retained intron. D-F) DAVID GO analysis to identify biological processes that are enriched for genes that are downregulation (D) or upregulation (E), and molecular functions (F) that are enriched in transcripts that are alternatively spliced in EDL muscles after Rbfox knockout. G) RNA-seq tracks showing Mef2a in control and Rbfox DKO muscles. Inclusion of β exon is dependent on Rbfox proteins. H) RT-PCR showing reduced PSI for Mef2a β exon in DKO muscle when compared to control muscle.

**Figure S4.**
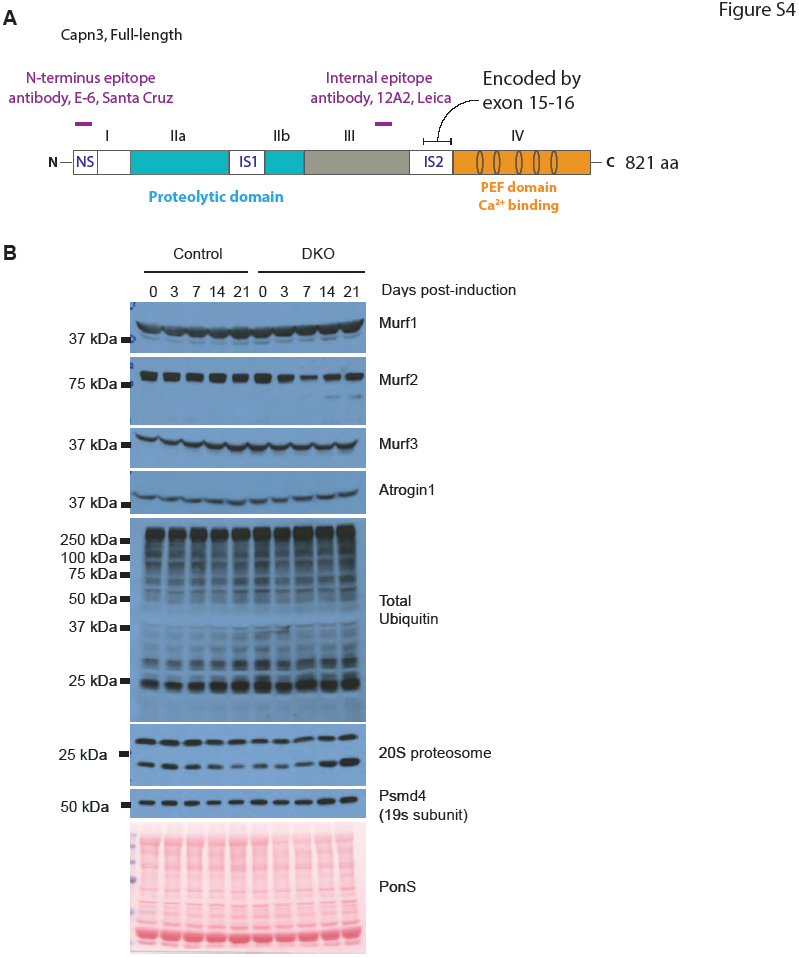
Altered proteostasis in Rbfox knockout muscles. A) Schematic showing four domains of Calpain 3 protein which are also present in Calpain family members. There are three unique Calpain3-specific insert sequences: N-terminus insert sequence (NS), Insert Sequence 1 (IS1), and 2 (IS2) are also indicated. Note that majority of the IS2 sequence is encoded by exons 15–16. Also indicates representative regions that are recognized by the two antibodies used in this study, the N-terminus (E-6, Santa Cruz) and the internal epitope (NCL-CALP-12A2, Leica) antibodies. B). Western blots using antibodies against indicated proteins in uninduced (0 day) and 3,7,14, and 21 days induced TA muscles from control and DKO animals.

## References

Bassel-Duby, R., and Olson, E.N. (2006). Signaling pathways in skeletal muscle remodeling. Annual review of biochemistry 75, 19–37.

Beckmann, J.S., and Spencer, M. (2008). Calpain 3, the “gatekeeper” of proper sarcomere assembly, turnover and maintenance. Neuromuscul Disord 18, 913–921.

Bell, R.A., Al-Khalaf, M., and Megeney, L.A. (2016a). The beneficial role of proteolysis in skeletal muscle growth and stress adaptation. Skeletal muscle 6, 16.

Bell, R.A., Al-Khalaf, M., and Megeney, L.A. (2016b). Erratum to: The beneficial role of proteolysis in skeletal muscle growth and stress adaptation. Skeletal muscle 6, 19.

Bodine, S.C., and Baehr, L.M. (2014). Skeletal muscle atrophy and the E3 ubiquitin ligases MuRF1 and MAFbx/atrogin-1. American journal of physiology Endocrinology and metabolism 307, E469–484.

Bodine, S.C., Latres, E., Baumhueter, S., Lai, V.K., Nunez, L., Clarke, B.A., Poueymirou, W.T., Panaro, F.J., Na, E., Dharmarajan, K., et al. (2001). Identification of ubiquitin ligases required for skeletal muscle atrophy. Science 294, 1704–1708.

Bowen, T.S., Schuler, G., and Adams, V. (2015). Skeletal muscle wasting in cachexia and sarcopenia: molecular pathophysiology and impact of exercise training. Journal of cachexia, sarcopenia and muscle 6, 197–207.

Charton, K., Suel, L., Henriques, S.F., Moussu, J.P., Bovolenta, M., Taillepierre, M., Becker, C., Lipson, K., and Richard, I. (2016). Exploiting the CRISPR/Cas9 system to study alternative splicing in vivo: application to titin. Hum Mol Genet 25, 4518–4532.

Comai, G., and Tajbakhsh, S. (2014). Molecular and cellular regulation of skeletal myogenesis. Current topics in developmental biology 110, 1–73.

Crossland, R.F., Balasa, A., Ramakrishnan, R., Mahadevan, S.K., Fiorotto, M.L., and Van den Veyver, I.B. (2017). Chronic Maternal Low-Protein Diet in Mice Affects Anxiety, Night-Time Energy Expenditure and Sleep Patterns, but Not Circadian Rhythm in Male Offspring. PLoS One 12, e0170127.

Fanin, M., Nascimbeni, A.C., Fulizio, L., Trevisan, C.P., Meznaric-Petrusa, M., and Angelini, C. (2003). Loss of calpain-3 autocatalytic activity in LGMD2A patients with normal protein expression. Am J Pathol 163, 1929–1936.

Gallagher, T.L., Arribere, J.A., Geurts, P.A., Exner, C.R., McDonald, K.L., Dill, K.K., Marr, H.L., Adkar, S.S., Garnett, A.T., Amacher, S.L., et al. (2011). Rbfox-regulated alternative splicing is critical for zebrafish cardiac and skeletal muscle functions. Dev Biol 359, 251–261.

Gehman, L.T., Meera, P., Stoilov, P., Shiue, L., O’Brien, J.E., Meisler, M.H., Ares, M., Jr., Otis, T.S., and Black, D.L. (2012). The splicing regulator Rbfox2 is required for both cerebellar development and mature motor function. Genes Dev 26, 445–460.

Gehman, L.T., Stoilov, P., Maguire, J., Damianov, A., Lin, C.H., Shiue, L., Ares, M., Jr., Mody, I., and Black, D.L. (2011). The splicing regulator Rbfox1 (A2BP1) controls neuronal excitation in the mammalian brain. Nat Genet 43, 706–711.

Goodman, C.A., and Hornberger, T.A. (2013). Measuring protein synthesis with SUnSET: a valid alternative to traditional techniques? Exercise and sport sciences reviews 41, 107–115.

Huang da, W., Sherman, B.T., and Lempicki, R.A. (2009). Systematic and integrative analysis of large gene lists using DAVID bioinformatics resources. Nat Protoc 4, 44–57.

Huang, J., and Forsberg, N.E. (1998). Role of calpain in skeletal-muscle protein degradation. Proc Natl Acad Sci U S A 95, 12100–12105.

James, H.A., O’Neill, B.T., and Nair, K.S. (2017). Insulin Regulation of Proteostasis and Clinical Implications. Cell Metab.

Jin, Y., Suzuki, H., Maegawa, S., Endo, H., Sugano, S., Hashimoto, K., Yasuda, K., and Inoue, K. (2003). A vertebrate RNA-binding protein Fox-1 regulates tissue-specific splicing via the pentanucleotide GCAUG. EMBO J 22, 905–912.

Ju, J.S., Varadhachary, A.S., Miller, S.E., and Weihl, C.C. (2010). Quantitation of “autophagic flux” in mature skeletal muscle. Autophagy 6, 929–935.

Kalsotra, A., and Cooper, T.A. (2011). Functional consequences of developmentally regulated alternative splicing. Nature reviews Genetics 12, 715–729.

Katz, Y., Wang, E.T., Airoldi, E.M., and Burge, C.B. (2010). Analysis and design of RNA sequencing experiments for identifying isoform regulation. Nature methods 7, 1009–1015.

Kramerova, I., Kudryashova, E., Tidball, J.G., and Spencer, M.J. (2004). Null mutation of calpain 3 (p94) in mice causes abnormal sarcomere formation in vivo and in vitro. Hum Mol Genet 13, 1373–1388.

Kuroyanagi, H., Ohno, G., Mitani, S., and Hagiwara, M. (2007). The Fox-1 family and SUP-12 coordinately regulate tissue-specific alternative splicing in vivo. Mol Cell Biol 27, 8612–8621.

Kwon, A.T., Arenillas, D.J., Worsley Hunt, R., and Wasserman, W.W. (2012). oPOSSUM-3: advanced analysis of regulatory motif over-representation across genes or ChIP-Seq datasets. G3 2, 987–1002.

Lecker, S.H., Goldberg, A.L., and Mitch, W.E. (2006). Protein degradation by the ubiquitin-proteasome pathway in normal and disease states. Journal of the American Society of Nephrology: JASN 17, 1807–1819.

Masiero, E., Agatea, L., Mammucari, C., Blaauw, B., Loro, E., Komatsu, M., Metzger, D., Reggiani, C., Schiaffino, S., and Sandri, M. (2009). Autophagy is required to maintain muscle mass. Cell Metab 10, 507–515.

Masiero, E., and Sandri, M. (2010). Autophagy inhibition induces atrophy and myopathy in adult skeletal muscles. Autophagy 6, 307–309.

McLeod, M., Breen, L., Hamilton, D.L., and Philp, A. (2016). Live strong and prosper: the importance of skeletal muscle strength for healthy ageing. Biogerontology 17, 497–510.

Nakajima, E., David, L.L., Bystrom, C., Shearer, T.R., and Azuma, M. (2006). Calpain-specific proteolysis in primate retina: Contribution of calpains in cell death. Investigative ophthalmology & visual science 47, 5469–5475.

Navone, F., Genevini, P., and Borgese, N. (2015). Autophagy and Neurodegeneration: Insights from a Cultured Cell Model of ALS. Cells 4, 354–386.

Nishino, I., Fu, J., Tanji, K., Yamada, T., Shimojo, S., Koori, T., Mora, M., Riggs, J.E., Oh, S.J., Koga, Y., et al. (2000). Primary LAMP-2 deficiency causes X-linked vacuolar cardiomyopathy and myopathy (Danon disease). Nature 406, 906–910.

Ono, Y., Kakinuma, K., Torii, F., Irie, A., Nakagawa, K., Labeit, S., Abe, K., Suzuki, K., and Sorimachi, H. (2004). Possible regulation of the conventional calpain system by skeletal muscle-specific calpain, p94/calpain 3. J Biol Chem 279, 2761–2771.

Ono, Y., Shindo, M., Doi, N., Kitamura, F., Gregorio, C.C., and Sorimachi, H. (2014). The N-and C-terminal autolytic fragments of CAPN3/p94/calpain-3 restore proteolytic activity by intermolecular complementation. Proc Natl Acad Sci U S A 111, E5527–5536.

Ono, Y., and Sorimachi, H. (2012). Calpains: an elaborate proteolytic system. Biochim Biophys Acta 1824, 224–236.

Pedrotti, S., Giudice, J., Dagnino-Acosta, A., Knoblauch, M., Singh, R.K., Hanna, A., Mo, Q., Hicks, J., Hamilton, S., and Cooper, T.A. (2015). The RNA-binding protein Rbfox1 regulates splicing required for skeletal muscle structure and function. Hum Mol Genet.

Piatkov, K.I., Oh, J.H., Liu, Y., and Varshavsky, A. (2014). Calpain-generated natural protein fragments as short-lived substrates of the N-end rule pathway. Proc Natl Acad Sci U S A 111, E817–826.

Potthoff, M.J., and Olson, E.N. (2007). MEF2: a central regulator of diverse developmental programs. Development 134, 4131–4140.

Raben, N., Hill, V., Shea, L., Takikita, S., Baum, R., Mizushima, N., Ralston, E., and Plotz, P. (2008). Suppression of autophagy in skeletal muscle uncovers the accumulation of ubiquitinated proteins and their potential role in muscle damage in Pompe disease. Hum Mol Genet 17, 3897–3908.

Rai, M., and Demontis, F. (2016). Systemic Nutrient and Stress Signaling via Myokines and Myometabolites. Annu Rev Physiol 78, 85–107.

Raj, B., and Blencowe, B.J. (2015). Alternative Splicing in the Mammalian Nervous System: Recent Insights into Mechanisms and Functional Roles. Neuron 87, 14–27.

Rao, P., and Monks, D.A. (2009). A tetracycline-inducible and skeletal muscle-specific Cre recombinase transgenic mouse. Developmental neurobiology 69, 401–406.

Ribas, V., Drew, B.G., Zhou, Z., Phun, J., Kalajian, N.Y., Soleymani, T., Daraei, P., Widjaja, K., Wanagat, J., de Aguiar Vallim, T.Q., et al. (2016). Skeletal muscle action of estrogen receptor alpha is critical for the maintenance of mitochondrial function and metabolic homeostasis in females. Sci Transl Med 8, 334ra354.

Richard, I., Broux, O., Allamand, V., Fougerousse, F., Chiannilkulchai, N., Bourg, N., Brenguier, L., Devaud, C., Pasturaud, P., Roudaut, C., et al. (1995). Mutations in the proteolytic enzyme calpain 3 cause limb-girdle muscular dystrophy type 2A. Cell 81, 27–40.

Robinson, M.D., McCarthy, D.J., and Smyth, G.K. (2010). edgeR: a Bioconductor package for differential expression analysis of digital gene expression data. Bioinformatics 26, 139–140.

Runfola, V., Sebastian, S., Dilworth, F.J., and Gabellini, D. (2015). Rbfox proteins regulate tissue-specific alternative splicing of Mef2D required for muscle differentiation. J Cell Sci 128, 631–637.

Saha, P.K., Reddy, V.T., Konopleva, M., Andreeff, M., and Chan, L. (2010). The triterpenoid 2-cyano-3,12-dioxooleana-1,9-dien-28-oic-acid methyl ester has potent anti-diabetic effects in diet-induced diabetic mice and Lepr(db/db) mice. J Biol Chem 285, 40581–40592.

Sandri, M., Coletto, L., Grumati, P., and Bonaldo, P. (2013). Misregulation of autophagy and protein degradation systems in myopathies and muscular dystrophies. J Cell Sci 126, 5325–5333.

Schiaffino, S., Dyar, K.A., Ciciliot, S., Blaauw, B., and Sandri, M. (2013). Mechanisms regulating skeletal muscle growth and atrophy. Febs J 280, 4294–4314.

Schmidt, E.K., Clavarino, G., Ceppi, M., and Pierre, P. (2009). SUnSET, a nonradioactive method to monitor protein synthesis. Nature methods 6, 275–277.

Sebastian, S., Faralli, H., Yao, Z., Rakopoulos, P., Palii, C., Cao, Y., Singh, K., Liu, Q.C., Chu, A., Aziz, A., et al. (2013). Tissue-specific splicing of a ubiquitously expressed transcription factor is essential for muscle differentiation. Genes Dev 27, 1247–1259.

Singh, R.K., Xia, Z., Bland, C.S., Kalsotra, A., Scavuzzo, M.A., Curk, T., Ule, J., Li, W., and Cooper, T.A. (2014). Rbfox2-Coordinated Alternative Splicing of Mef2d and Rock2 Controls Myoblast Fusion during Myogenesis. Mol Cell 55, 592–603.

Smith, H.L., Li, W., and Cheetham, M.E. (2015). Molecular chaperones and neuronal proteostasis. Seminars in cell & developmental biology 40, 142–152.

Spencer, M.J., Guyon, J.R., Sorimachi, H., Potts, A., Richard, I., Herasse, M., Chamberlain, J., Dalkilic, I., Kunkel, L.M., and Beckmann, J.S. (2002). Stable expression of calpain 3 from a muscle transgene in vivo: immature muscle in transgenic mice suggests a role for calpain 3 in muscle maturation. Proc Natl Acad Sci U S A 99, 8874–8879.

Sulistio, Y.A., and Heese, K. (2016). The Ubiquitin-Proteasome System and Molecular Chaperone Deregulation in Alzheimer’s Disease. Molecular neurobiology 53, 905–931.

Takamure, M., Murata, K.Y., Tamada, Y., Azuma, M., and Ueno, S. (2005). Calpain-dependent alpha-fodrin cleavage at the sarcolemma in muscle diseases. Muscle Nerve 32, 303–309.

Tanaka, Y., Guhde, G., Suter, A., Eskelinen, E.L., Hartmann, D., Lullmann-Rauch, R., Janssen, P.M., Blanz, J., von Figura, K., and Saftig, P. (2000). Accumulation of autophagic vacuoles and cardiomyopathy in LAMP-2-deficient mice. Nature 406, 902–906.

Taveau, M., Bourg, N., Sillon, G., Roudaut, C., Bartoli, M., and Richard, I. (2003). Calpain 3 is activated through autolysis within the active site and lyses sarcomeric and sarcolemmal components. Mol Cell Biol 23, 9127–9135.

Trapnell, C., Williams, B.A., Pertea, G., Mortazavi, A., Kwan, G., van Baren, M.J., Salzberg, S.L., Wold, B.J., and Pachter, L. (2010). Transcript assembly and quantification by RNA-Seq reveals unannotated transcripts and isoform switching during cell differentiation. Nat Biotechnol 28, 511–515.

Venables, J.P., Vignal, E., Baghdiguian, S., Fort, P., and Tazi, J. (2012). Tissue-specific alternative splicing of Tak1 is conserved in deuterostomes. Molecular biology and evolution 29, 261–269.

Wang, E.T., Sandberg, R., Luo, S., Khrebtukova, I., Zhang, L., Mayr, C., Kingsmore, S.F., Schroth, G.P., and Burge, C.B. (2008). Alternative isoform regulation in human tissue transcriptomes. Nature 456, 470–476.

Yang, X., Coulombe-Huntington, J., Kang, S., Sheynkman, G.M., Hao, T., Richardson, A., Sun, S., Yang, F., Shen, Y.A., Murray, R.R., et al. (2016). Widespread Expansion of Protein Interaction Capabilities by Alternative Splicing. Cell 164, 805–817.

Yin, H., Price, F., and Rudnicki, M.A. (2013). Satellite cells and the muscle stem cell niche. Physiol Rev 93, 23–67.

Zhang, Y., and Bhavnani, B.R. (2006). Glutamate-induced apoptosis in neuronal cells is mediated via caspase-dependent and independent mechanisms involving calpain and caspase-3 proteases as well as apoptosis inducing factor (AIF) and this process is inhibited by equine estrogens. BMC neuroscience 7, 49.

